# Selective neuroimmune modulation by type I interferon drives neuropathology and neurologic dysfunction following traumatic brain injury

**DOI:** 10.1101/2023.06.06.543774

**Authors:** Brittany P. Todd, Zili Luo, Noah Gilkes, Michael S. Chimenti, Zeru Peterson, Madison Mix, John T. Harty, Thomas Nickl-Jockschat, Polly J. Ferguson, Alexander G. Bassuk, Elizabeth A. Newell

**Affiliations:** Medical Scientist Training Program, University of Iowa, Iowa City, IA, USA; Department of Pediatrics, University of Iowa, Iowa City, IA, USA; Iowa Institute of Human Genetics, Bioinformatics Division, University of Iowa, Iowa City, IA, USA; Department of Neuroscience and Pharmacology, University of Iowa, Iowa City, IA, USA; Department of Pathology and Interdisciplinary Graduate Program in Immunology, University of Iowa, IA, USA; Department of Psychiatry, University of Iowa, Iowa City, IA, USA

## Abstract

Accumulating evidence suggests that type I interferon (IFN-I) signaling is a key contributor to immune cell-mediated neuropathology in neurodegenerative diseases. Recently, we demonstrated a robust upregulation of type I interferon-stimulated genes in microglia and astrocytes following experimental traumatic brain injury (TBI). The specific molecular and cellular mechanisms by which IFN-I signaling impacts the neuroimmune response and neuropathology following TBI remains unknown. Using the lateral fluid percussion injury model (FPI) in adult male mice, we demonstrated that IFN α/β receptor (IFNAR) deficiency resulted in selective and sustained blockade of type I interferon-stimulated genes following TBI as well as decreased microgliosis and monocyte infiltration. Phenotypic alteration of reactive microglia also occurred with diminished expression of molecules needed for MHC class I antigen processing and presentation following TBI. This was associated with decreased accumulation of cytotoxic T cells in the brain. The IFNAR-dependent modulation of the neuroimmune response was accompanied by protection from secondary neuronal death, white matter disruption, and neurobehavioral dysfunction. These data support further efforts to leverage the IFN-I pathway for novel, targeted therapy of TBI.

## Introduction

Traumatic brain injury (TBI) is a leading cause of death and disability through young adulthood; patients who survive their injury often develop chronic neurological disease (1–3). Despite the high burden of TBI, neuroprotective therapies do not exist. The current lack of treatment options stems from an incomplete understanding of the mechanisms that lead to ongoing neurologic dysfunction and degeneration after injury.

Following TBI, cellular damage initiates acute immune reactivity including activation of resident glial cells, infiltration of peripheral leukocytes, and increased production of soluble inflammatory mediators (4). While the initial surge in immune activation facilitates debris clearance and regeneration, chronic dysregulated immune cell reactivity may contribute to progressive neurodegeneration and pathology. Growing evidence suggests that microglia are key cellular mediators of chronic neurologic dysfunction following TBI, with microglia depletion resulting in decreased inflammation and reduced neuropathology (5–7). The specific mechanisms that promote chronic microglial and peripheral immune cell activation following traumatic brain injury remain unclear.

One potential mechanism underlying sustained immune activation is type I interferon (IFN-I) signaling through the interferon-α/β receptor (IFNAR). IFNAR is encoded by two genes, IFNAR1 and IFNAR2, both are essential for receptor function. Type I IFNs are proteins that regulate the recruitment and effector functions of immune cells (8). Our lab has demonstrated that following TBI, the microglial transcriptome is highly enriched for type I interferon-stimulated genes (ISGs) (9). Others have shown upregulation of ISGs in the cortex and hippocampus following experimental TBI (5, 10, 11). The robust upregulation of the IFN-I pathway at both the tissue and immune cell level suggests that IFN-I signaling is a potential instigator of sustained, dysregulated immune response and a source of neuropathology. Initial studies of IFN-I deficiency after TBI demonstrate reduced acute inflammatory gene expression and ISG expression that was associated with improved neurologic function (10, 11). However, the prior studies are limited by their emphasis on acute timepoints after TBI and by their lack of mechanistic data on how specific immune cell types respond to TBI. The objective of the following study is to determine the effect of IFN-I signaling on subacute and chronic neuroimmune activation, neuropathology, and neurologic function following TBI.

## Results

### Activation of the type I interferon pathway is persistent and widespread in the brain after TBI

We had demonstrated that the transcriptional response of microglia is highly enriched for expression of genes in the type I interferon pathway (IFN-I) at seven days following TBI (9). We now sought to characterize the time course and spatial localization of IFN-I pathway upregulation following TBI. We selected a panel of interferon-stimulated genes (ISGs) that were upregulated in microglia seven days post-injury (DPI) including *Ifit3*, *Ifi204*, *Stat1*, *Irf7*, and *Axl* (9). Using quantitative real-time PCR (qPCR), we evaluated expression of these ISGs in regional brain tissue at 1 DPI through 21 DPI. We found that in both the ipsilateral perilesional cortex and hippocampus, ISGs were elevated by 1 DPI and several ISGs remained elevated at 21 DPI (**Figure 1, A and B**). We next performed dual RNAscope and immunohistochemistry (IHC) to assess spatial location and confirm the cell source of *Irf7*, a key transcriptional factor in IFN-I pathway activation. We found that at 7 DPI, reactive microglia were a substantial source of *Irf7* expression in widespread regions of the brain (**Figure 1, C-F**). These reactive microglia were present in the cortex and hippocampus as well as two areas known to be affected by traumatic axonal injury, the corpus callosum and thalamus.

**Figure 1:**
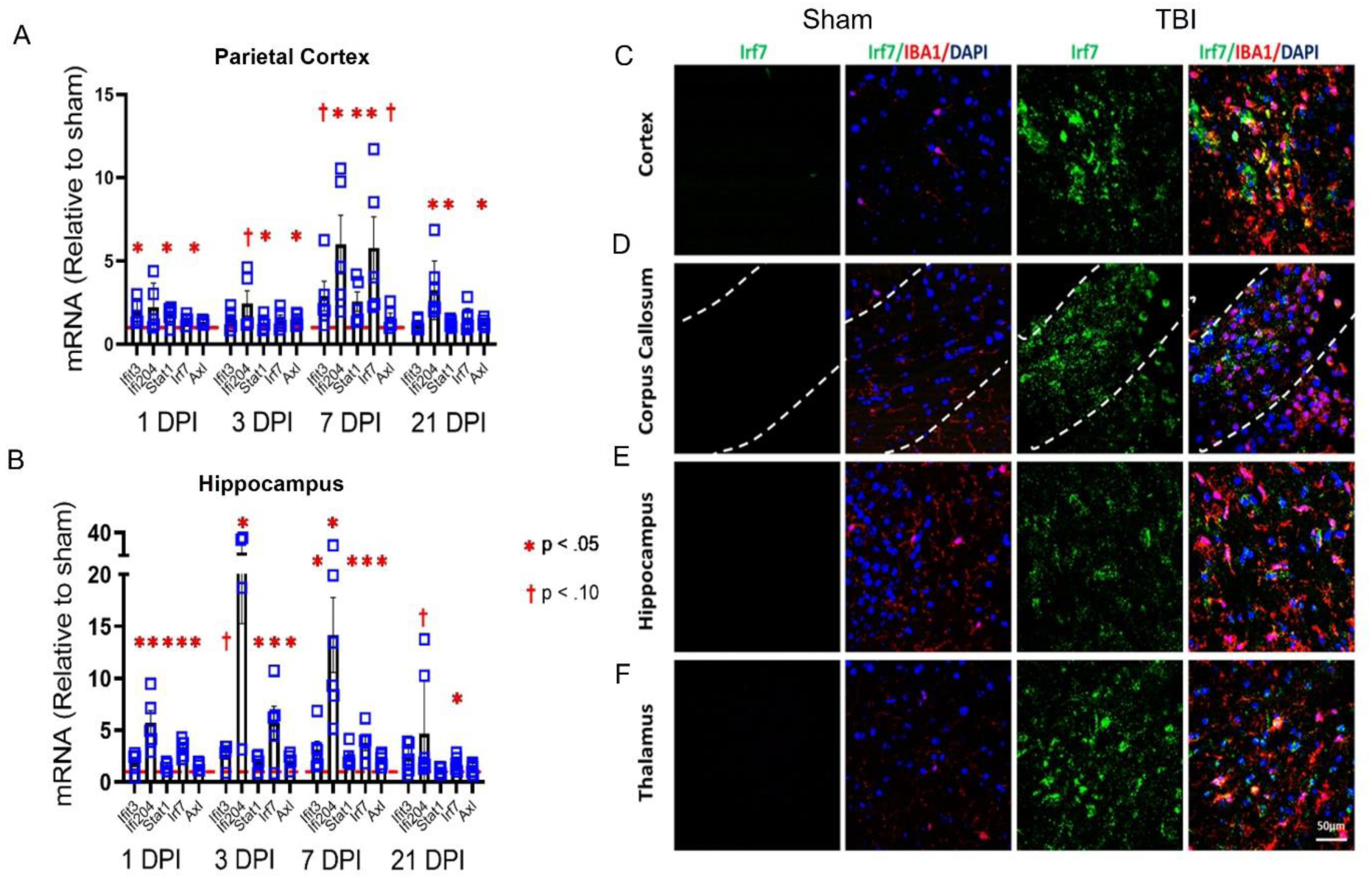
TBI results in rapid and sustained ISG expression in both focal and remote regions of the brain. ISG expression was assessed from ipsilateral cortex (**A**) or hippocampus (**B**) at various timepoints post injury by qPCR. Data are expressed as fold change relative to sham. Blue squares represent individual animals. Red dashed line indicates mean sham level of gene expression. N=5-6 mice/group. Significance determined by two-way ANOVA with Fisher’s LSD for multiple comparisons.*p<0.05 TBI compared to sham, ✝p<0.1 TBI compared to sham. (**C-F**) Representative confocal images demonstrating TBI-induced, microglial *Irf7* expression. At 7 days post-TBI, *Irf7* co-localized with the microglial marker, IBA1 (Scale bar, 50 um).

### IFNAR deficiency modulates neuroinflammatory gene expression following TBI

Given the pleiotropic and immune-modulating functions of IFN-I, we next sought to assess the impact of IFNAR deficiency on neuroinflammatory gene expression following TBI (12, 13). We used IFNAR1 KO mice which resulted in total loss of IFNAR receptor function. The NanoString neuroinflammation panel was used to evaluate the expression of 770 genes that are involved in primary immune function/inflammatory processes. TBI was induced in wild-type and IFN-I signalling-deficient mice by lateral fluid percussion injury (FPI). We used pairwise comparison of WT TBI vs. IFNAR KO TBI to determine differentially expressed genes (DEGs). There were 10 genes (*Irf7, Rsad2, Slfn8, Oas1g, Ddx58, Ly6a, Ifih1, Ifitm3, Zbp1, Cxcl10*) that were significantly decreased in IFNAR KO TBI compared to WT TBI (**Figure 2A)**. It is important to note that there were no significant DEGs when comparing WT and IFNAR KO uninjured controls. All 10 DEGs have been described as interferon-stimulated genes (ISGs). These genes have many functions including positive regulation of anti-viral immune activation (*Irf7*), nucleic acid sensing (*Ddx58, Ifih1, Oas1g, Zbp1*), T cell activation and differentiation (*Rsad2, Slfn8*), and chemoattraction (*Cxcl10*). However, broad suppression of the inflammatory response to TBI did not occur: IFNAR KO mice still had 366 DEGs in response to TBI, and these included many inflammation-related genes that have been described as upregulated following TBI (**Supplemental Figure 1, A and B, Table 1**). In conclusion, we found that IFNAR deficiency resulted in the modulation of a specific subset of inflammatory genes following TBI.

**Figure 2:**
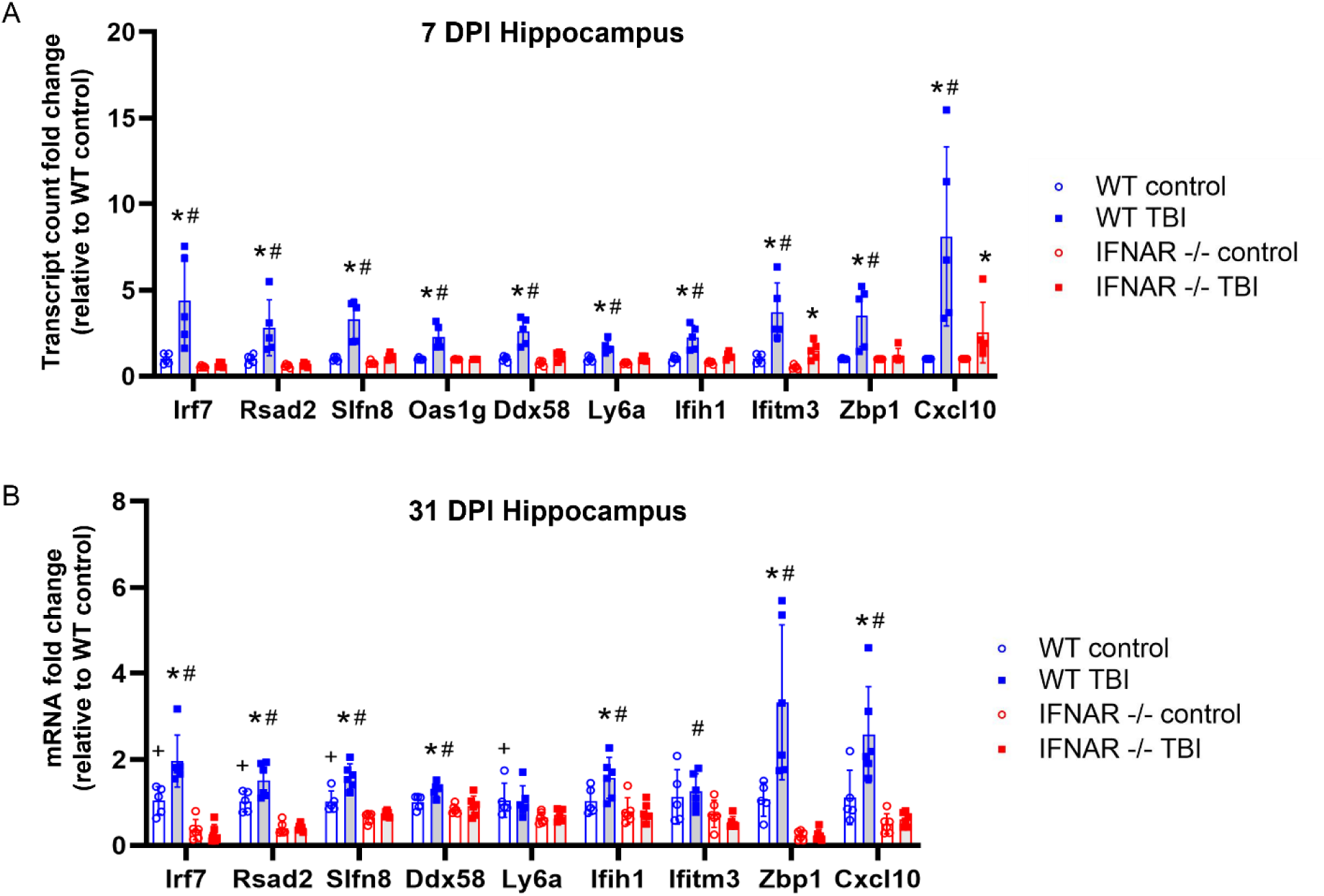
IFNAR deficiency results in decreased ISG expression following TBI. mRNA was isolated from hippocampal tissue at 7 and 31 DPI. (**A**) Gene expression at 7 DPI was evaluated using the NanoString Neuroinflammation panel. mRNA counts were compared in WT and IFNAR KO control vs WT and IFNAR KO TBI mice. Data are expressed as fold change in mRNA expression relative to WT control. Significance was determined by DESeq2 analysis. (**B**) Hippocampal gene expression at 31 DPI was evaluated by quantitative PCR. Significance was determined by two-way ANOVA with Fisher’s LSD for multiple comparisons. *p<0.05 TBI compared to matched genotype control, #p<0.05 WT TBI vs IFNAR KO TBI, +p<0.05 WT control vs IFNAR control.

To determine whether IFNAR signaling also impacts the chronic transcriptional response to TBI, we collected hippocampi from WT and IFNAR KO mice at thirty-one days post-injury (31 DPI) and performed qPCR (**Figure 2B**). We found that expression of seven of the ten DEGs identified at 7 DPI remained elevated in WT TBI at 31 DPI compared to WT controls. This increased expression was IFNAR-dependent, as IFNAR KO TBI mice had significantly less expression of all seven genes compared to WT TBI mice (*Irf7, Rsad2, Slfn8, Ddx58, Ifih1, Zbp1, Cxcl10;* p<0.001). Overall, IFN-I pathway genes remained elevated chronically post-TBI and this was prevented by IFNAR deficiency. The specific functions of these IFN-I pathway genes may provide clues as to how IFNAR deficiency alters outcomes following TBI.

One of the TBI-induced genes modulated by IFNAR deficiency, *Cxcl10*, is well recognized for its involvement in a variety of neurologic diseases (14). CXCL10 is a potent chemokine known to induce microglial reactivity and chemotaxis as well as recruitment of peripheral leukocytes to the central nervous system (CNS). To test the validity of our NanoString results and assess the spatial distribution of *Cxcl10*, we performed RNAscope in situ hybridization from brain tissue sections at 7 DPI. In accordance with our NanoString results, we observed upregulation of *Cxcl10* in WT TBI mice with minimal *Cxcl10* expression in IFNAR KO TBI mice as well as in both WT and IFNAR KO control mice. WT TBI animals had increased *Cxcl10* expression in the perilesional cortex and ipsilesional corpus callosum, hippocampus, and thalamus (**Figure 3, B-E**). Interestingly, the thalamus displayed the most robust *Cxcl10* expression compared to other regions after WT TBI. Although IFNAR KO TBI mice did have detectable *Cxcl10* expression in the perilesional cortex, corpus callosum, and hippocampus (**Figure 3, G-I**), it was minimal compared to *Cxcl10* expression in WT TBI mice. Additionally, *Cxcl10* staining was notably absent in the ipsilesional thalamus of IFNAR KO TBI subjects **(Figure 3J**). Overall, *Cxcl10* expression is increased throughout the brain 7 days following TBI. This expression is substantially reduced in IFNAR-deficient mice following injury.

**Figure 3:**
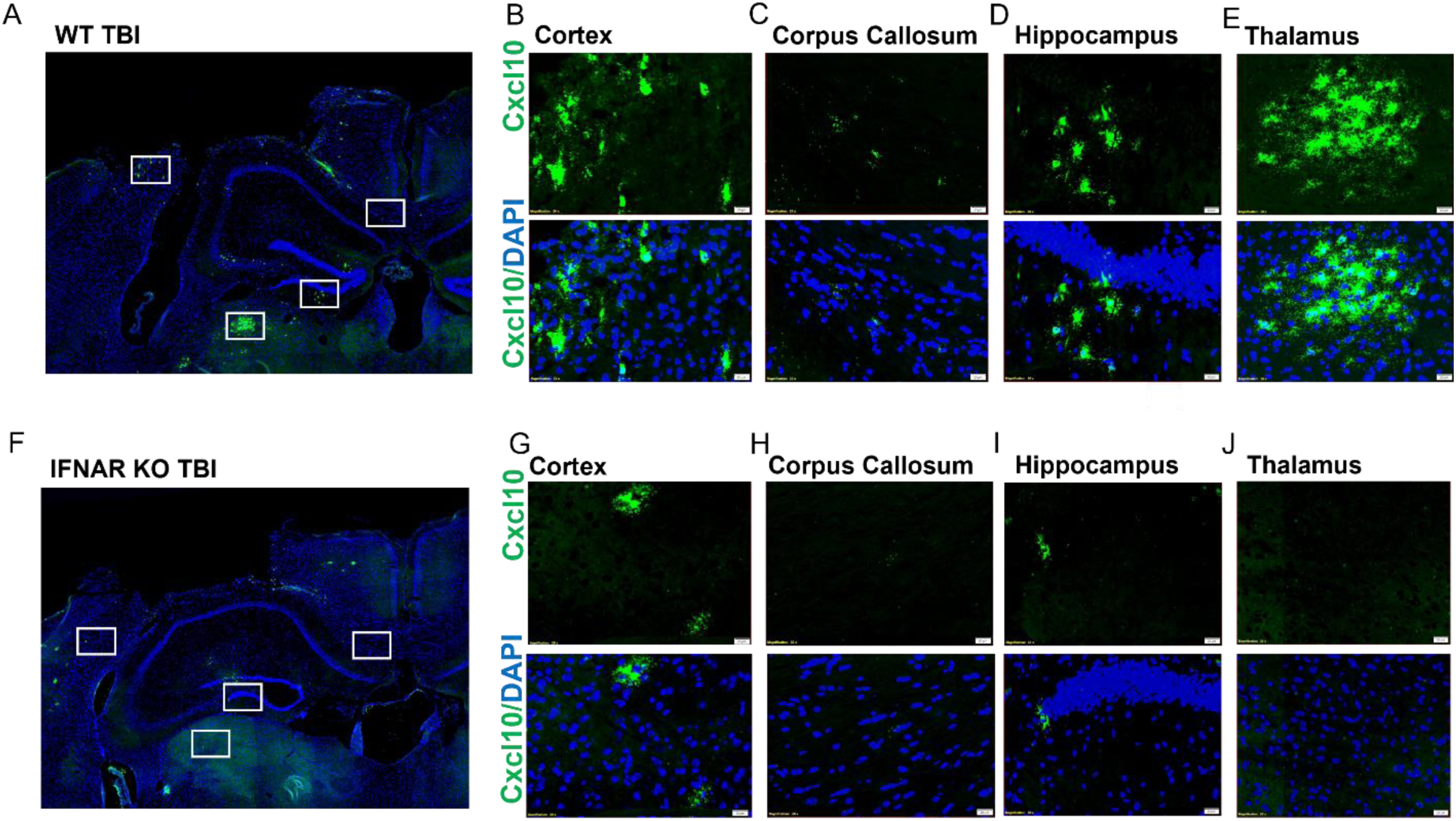
IFNAR deficiency decreases *Cxcl10* gene expression throughout the ipsilateral forebrain. Evaluation of TBI-induced *Cxcl10* expression using RNAscope in-situ hybridization: *Cxcl10* (green), DAPI (blue). 7 days post injury, WT mice (**A**) had greater *Cxcl10* expression than IFNAR KO mice (**F**) in the ipsilateral cortex (**B** and **G**), corpus callosum (**C** and **H**), hippocampus (**D** and **I**), and thalamus (**E** and **J**). Scale bar=20 um. WT and IFNAR KO control sections are not shown. Representative images from n=4-5 mice/group.

### IFNAR deficiency reduces microgliosis following TBI

After injury, reactive microglia migrate to sites of injury where they serve many functions including phagocytosis of debris, production of cytokines and chemokines, and antigen presentation (15). Given that microglia both produce and respond to type I interferons, we were interested in the effect of IFNAR deficiency on microgliosis following brain injury. To assess microglial reactivity and accumulation, we used IHC to stain for IBA1, a marker of microglia and macrophages, and calculated proportional area of IBA1 staining 31 days following TBI. As expected, WT mice had increased IBA1 staining in the thalamus 31 days following TBI (**Figure 4**). IFNAR KO TBI mice had greater thalamic microglial staining compared to IFNAR KO controls, but this injury-induced response was significantly less than WT TBI mice (**Figure 4F**). Both WT TBI and IFNAR KO TBI mice showed no difference in hippocampal IBA1 staining compared to their respective uninjured controls (**Figure 4**). In summary, IFNAR deficiency decreased microglial accumulation 31 days after TBI.

**Figure 4:**
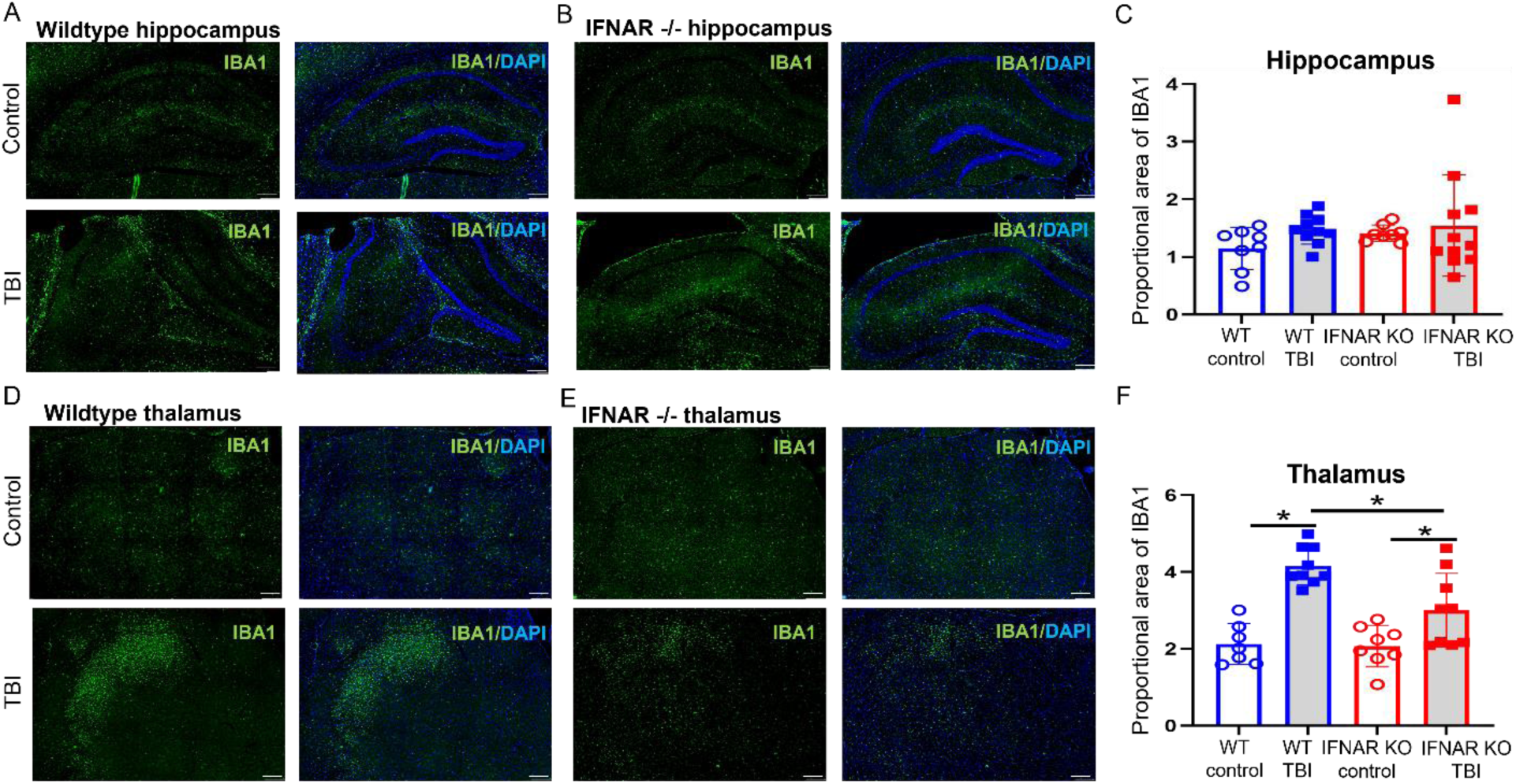
IFNAR deficiency reduces microgliosis following TBI. Microglial reactivity was visualized using IBA1 staining (green) 31 days following TBI. Representative IBA1 staining in the hippocampus (**A** and **B**) and thalamus (**D** and **E**) from injured WT and IFNAR KO mice. Scale bar=200 um. Quantification of % IBA1 area in thalamus (**C** and **F**). Error bars depict mean ± SEM. Statistical analysis performed by one-way ANOVA with Fisher’s LSD for multiple comparisons. *p≤0.05, n=7-9 mice/group.

### IFNAR deficiency reduces expression of tissue and microglial-specific MHC class I genes following TBI

In addition to evaluating microglial accumulation, we were interested in assessing the impact of IFNAR deficiency on the microglial phenotype following TBI. A key function of microglia is antigen presentation (16). We previously observed that several MHC class I molecules were upregulated in microglia following TBI (9). As many antigen-presentation and processing molecules are type I interferon-stimulated genes, we hypothesized that IFNAR deficiency would reduce MHC class I molecule expression in microglia following TBI. We first examined the impact of IFNAR deficiency on hippocampal expression of MHC class I genes, *H2-K1,* β*2m,* and *Tap1*, at 7 and 31 DPI (**Figure 5, A-C**). TBI resulted in increased hippocampal expression of *H2-K1,* β*2m,* and *Tap1* in WT mice at both timepoints. Conversely, TBI-induced upregulation of these MHC class I molecules was restricted in IFNAR KO mice. IFNAR KO TBI mice showed no increase in *H2-K1* expression at either time point compared to controls. IFNAR KO TBI mice had decreased expression of β*2m* compared to WT TBI mice at both timepoints. Lastly, while at 7 DPI, IFNAR KO TBI mice had a similar increase of *Tap1* expression as WT TBI mice, IFNAR KO mice showed a more rapid resolution of *Tap1* expression. By 31 DPI, *Tap1* expression was no longer elevated in IFNAR KO TBI compared to uninjured IFNAR KO controls.

**Figure 5:**
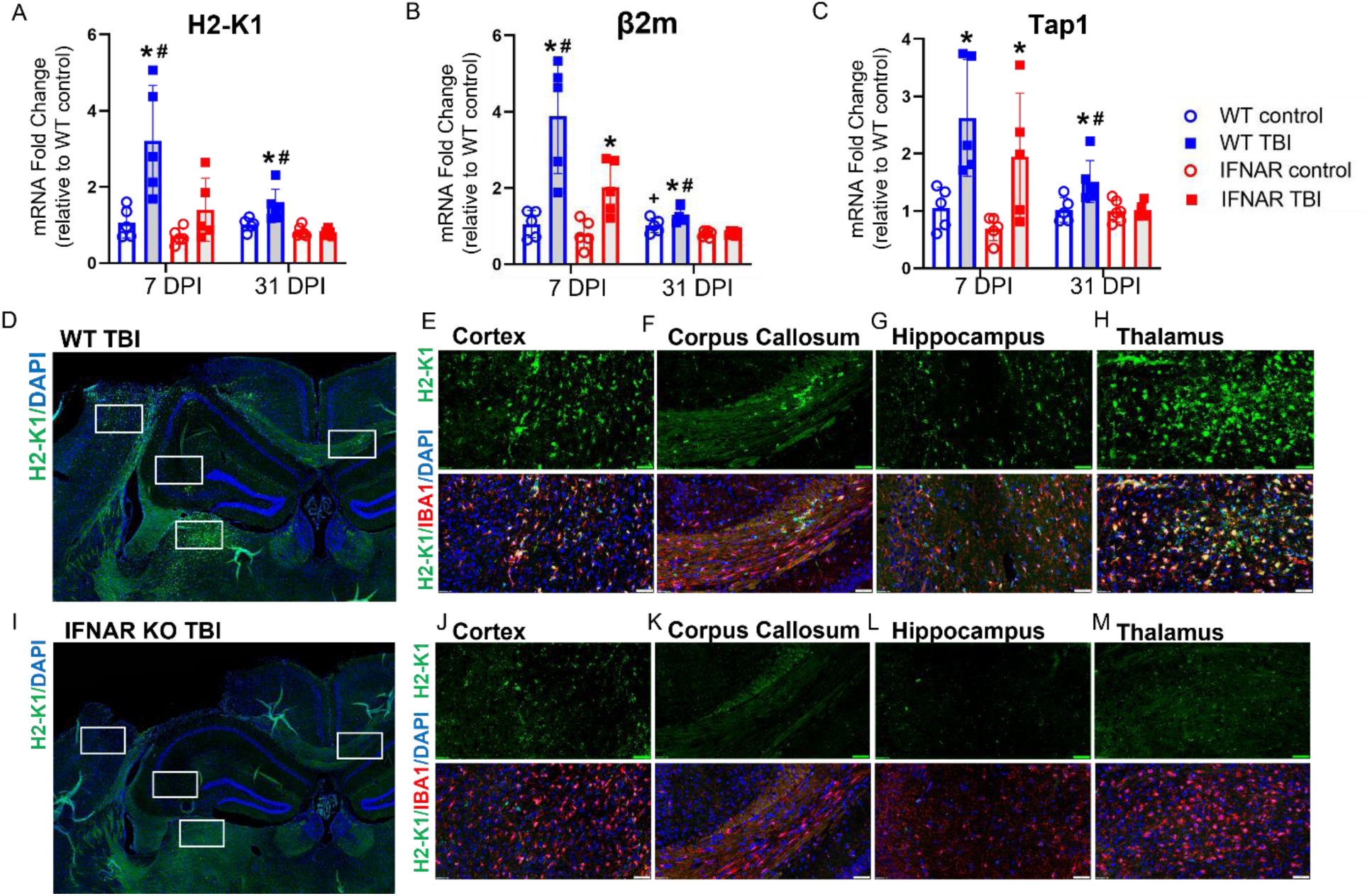
IFNAR deficiency reduces tissue and microglial MHC class I molecule expression at 7 and 31 days post-TBI. Expression of *H2-K1* (**A**) *β2m* (**B**) and *Tap1* (**C**) were evaluated by qPCR in ipsilateral hippocampus at 7 and 31 DPI in WT and IFNAR KO mice. Data are expressed as fold change in gene expression relative to WT control. Statistical analysis with two-way ANOVA with Fisher’s LSD for multiple comparisons. *p<0.05 for TBI compared to matched genotype control, # p<0.05 for WT TBI vs IFNAR KO TBI, n=5 mice/group. Evaluation of TBI-induced microglial *H2-K1* expression using dual in-situ hybridization and IHC: *H2-K1* (green), IBA1 (red), DAPI (blue). 7 DPI WT mice (**D**-**H**) had greater *H2-K1* expression and co-localization with a microglial marker, IBA1, than IFNAR KO mice (**I**-**M**) in all regions examined. Scale bar=50 um. WT and IFNAR KO control images are not shown. Representative images from n=4-5 mice/group.

Next, we combined fluorescent in situ hybridization with immunofluorescence to evaluate if the IFNAR-dependent reduction in MHC class I antigen processing and presentation molecules occurred specifically in reactive microglia. We used RNAscope to stain for *H2-K1* mRNA, and IBA1 immunostaining to identify microglia in tissue sections obtained from WT and IFNAR KO mice at 7 DPI. *H2-K1* expression was minimal in both WT and IFNAR KO control subjects. Following TBI in WT mice, *H2-K1* upregulation was seen in the ipsilesional cortex, corpus callosum, hippocampus, and thalamus (**Figure 5, D-H**) and colocalized with IBA1. The ipsilesional thalamus displayed the greatest injury-induced expression. In contrast, IFNAR KO TBI mice had minimal *H2-K1* staining compared to WT TBI animals. The perilesional cortex had the most *H2-K1* expression in the IFNAR KO subjects and similarly co-localized with IBA1 (**Figure 5, I-M**). Overall, IFNAR deficiency decreased microglial MHC class I molecule expression following TBI suggesting that IFNAR deficiency alters the phenotype of microglia and decreases their accumulation following TBI.

### IFNAR deficiency reduces monocyte populations following TBI

The IFNAR-mediated decrease in chemokine expression following TBI, suggested that IFNAR deficiency may impact the recruitment of peripheral immune cells to the CNS. To better understand how IFNAR deficiency alters the dynamics of immune cell populations in the CNS after TBI, we used flow cytometry to enumerate myeloid cell markers from whole brains at 3 and 10 DPI. Our gating strategy is shown in **Figure 6A**. Neither injury nor genotype significantly altered the number of microglia (**Figure 6B**). The total population of CD11b+, CD45+ high leukocytes was significantly increased at both 3 and 10 DPI in WT mice. IFNAR KO mice also showed an increase in CD11b+, CD45+ high leukocytes at 3 DPI with a trending increase at 10 DPI (**Figure 6C**). To evaluate the leukocyte identity, we stained for cell markers that demarcate neutrophils and monocytes. Neutrophils have previously been shown to rapidly infiltrate the brain after injury and decline in number in the days following injury (17). Our results were consistent with this understanding. Neutrophils were increased at 3 days post injury in WT TBI mice compared to controls. In contrast, IFNAR KO mice showed no increase in neutrophil counts at three or ten days post-injury (**Figure 6D**)

**Figure 6:**
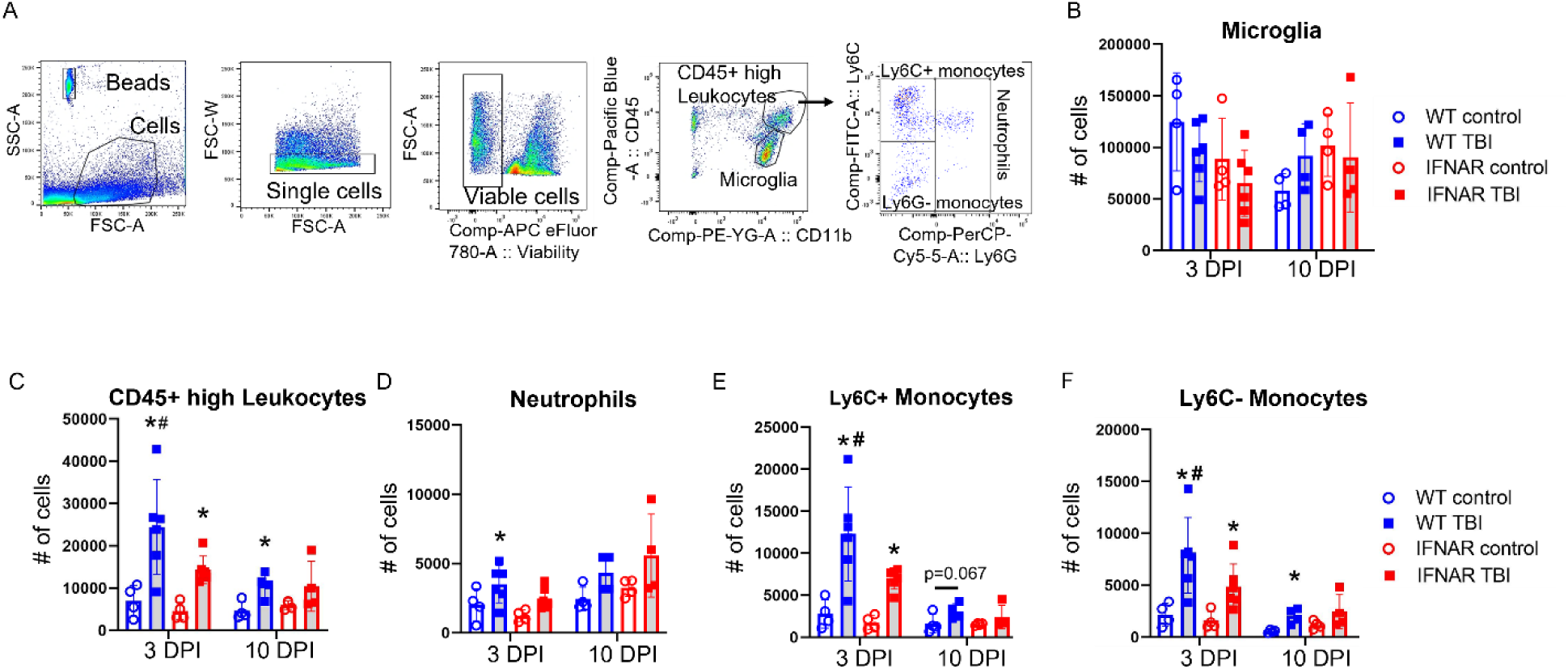
IFNAR deficiency reduces acute monocyte accumulation following TBI. Leukocyte cell accumulation was quantified using flow cytometry. (**A**) Representative flow cytometry gating strategy. (**B**) Total microglial numbers were not affected by genotype or brain injury. (**C**) TBI resulted in an increase in the total population of CD45+ high leukocytes. More specifically, the neutrophil population was increased acutely after WT TBI. (**D**) Ly6C+ and Ly6C-monocytes were increased at 3 and 10 days following WT TBI, while IFNAR KO mice showed significantly less monocyte accumulation after TBI (**E** and **F**). n=4 for control groups, n=4-6 for TBI groups. Error bars depict mean ± SEM. Statistical analysis performed by one-way ANOVA with Fisher’s LSD. * p<0.05 for control vs. TBI, # p<0.05 for WT TBI vs IFNAR KO TBI.

Monocytes also infiltrate the brain after injury, classically peaking in number at 3-4 days post injury, including both inflammatory (LY6C+) and patrolling (LY6C-) monocytes (17–19). In our model, WT TBI mice had increased LY6C+ and LY6C-monocytes at both time points. IFNAR KO TBI mice had significantly less LY6C+ and LY6C-monocytes than WT TBI mice at 3 DPI. At 10 DPI, IFNAR KO TBI were no different than genotypic uninjured controls. These results are evidence of decreased accumulation and hastened resolution of monocyte accumulation in IFNAR KO TBI compared to WT TBI mice (**Figure 6, E and F**).

### IFNAR deficiency reduces T cell accumulation following TBI

Decreased expression of the chemokine CXCL10 and of MHC class I presentation molecules, in tandem with reduced infiltration of monocyte populations following TBI, led us to hypothesize that IFNAR deficiency may also affect T cell accumulation following TBI. To investigate this question, we used flow cytometry to quantify the total number of CD4+ and CD8+ T cells in the whole brain at 3 and 10 DPI (**Figure 7A**). In agreement with prior studies (20–22), WT TBI mice had elevated CD4+ T cell counts at both 3 and 10 DPI compared to uninjured genotypic controls. However, this increase in CD4+ T cells was not seen in IFNAR KO TBI mice (**Figure 7B**). CD8+ T cells were also significantly increased in WT TBI mice at 3 and 10 DPI. In contrast, there was no TBI-induced increase in CD8+ T cells in IFNAR KO mice compared to IFNAR KO controls (**Figure 7C**). Overall, IFNAR deficiency reduced the accumulation of T cells in the brain following TBI, with the greatest effect on CD8+ T cells. To gain a better understanding of where CD8+ T cells are acting in the brain after injury, we stained for CD8α at 7 DPI. CD8α+ T cells were present in the perilesional cortex, and ipsilesional hippocampus, corpus callosum, and thalamus (**Figure 7, D-H**), suggesting that they are active in focal regions around the injury, white matter tracts, and subcortical structures such as the hippocampus and thalamus.

**Figure 7:**
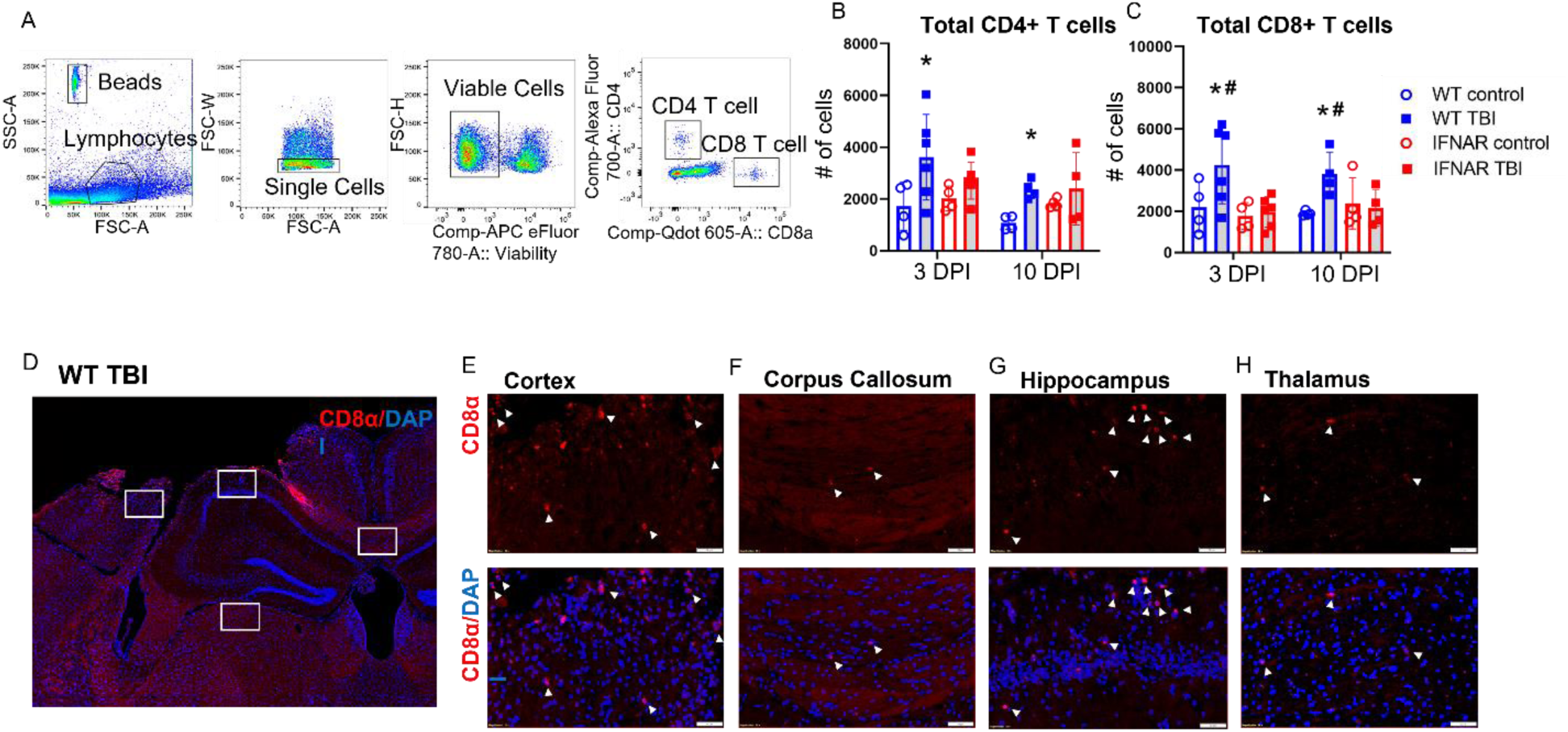
IFNAR deficiency reduces T cell accumulation following TBI. T cell accumulation was quantified using flow cytometry. (**A**) Representative flow cytometry gating strategy. TBI increased the total number of CD4+ T cells (**B**) and of CD8+ T cells (**C**) at 3 and 10 DPI days in WT mice; IFNAR KO mice showed no difference in T cell numbers after TBI compared to the IFNAR KO control, and a significant decrease in total CD8+ T cells compared to WT TBI at 3 and 10 DPI. n=4 for control groups, n=4-6 for TBI groups. Error bars depict mean ± SEM. Statistical analysis performed by two-way ANOVA with Fisher’s LSD. * p<0.05 for WT control vs. WT TBI, # p<0.05 for WT TBI vs IFNAR KO TBI. (**D**-**H**) Representative IHC for CD8α demonstrating localization of CD8+ T cells (arrowheads) in the cortex (**E**), corpus callosum (**F**), hippocampus (**G**), and thalamus (**H**) at 7 days following WT TBI. Scale bar=50 um. WT control sections not shown. n=4 mice/group.

### IFNAR deficiency reduces neuronal loss and white matter disruption following TBI

Neuronal death and degeneration are well documented after traumatic brain injury and may contribute to cognitive dysfunction (23–25). To determine whether IFNAR deficiency attenuates injury-induced neuronal loss, we stained for the neuronal marker, NeuN, in WT and IFNAR KO tissue sections obtained at 7 and 31 DPI. The thalamus was chosen as the region of interest in light of the IFNAR-dependent thalamic expression of *Cxcl10* and *H2-K1*, and the microglial accumulation observed in the thalamus. At both 7 and 31 DPI, WT TBI mice had significantly fewer thalamic neurons compared to uninjuredcontrols **(Figure 8).** IFNAR KO TBI mice showed no decrease in thalamic neuronal counts compared to uninjured controls at both timepoints and had significantly more thalamic neurons compared to WT TBI mice at 7 DPI **(Figure 8).** Overall, IFNAR deficiency attenuated thalamic neuronal loss at subacute and chronic timepoints following injury.

**Figure 8:**
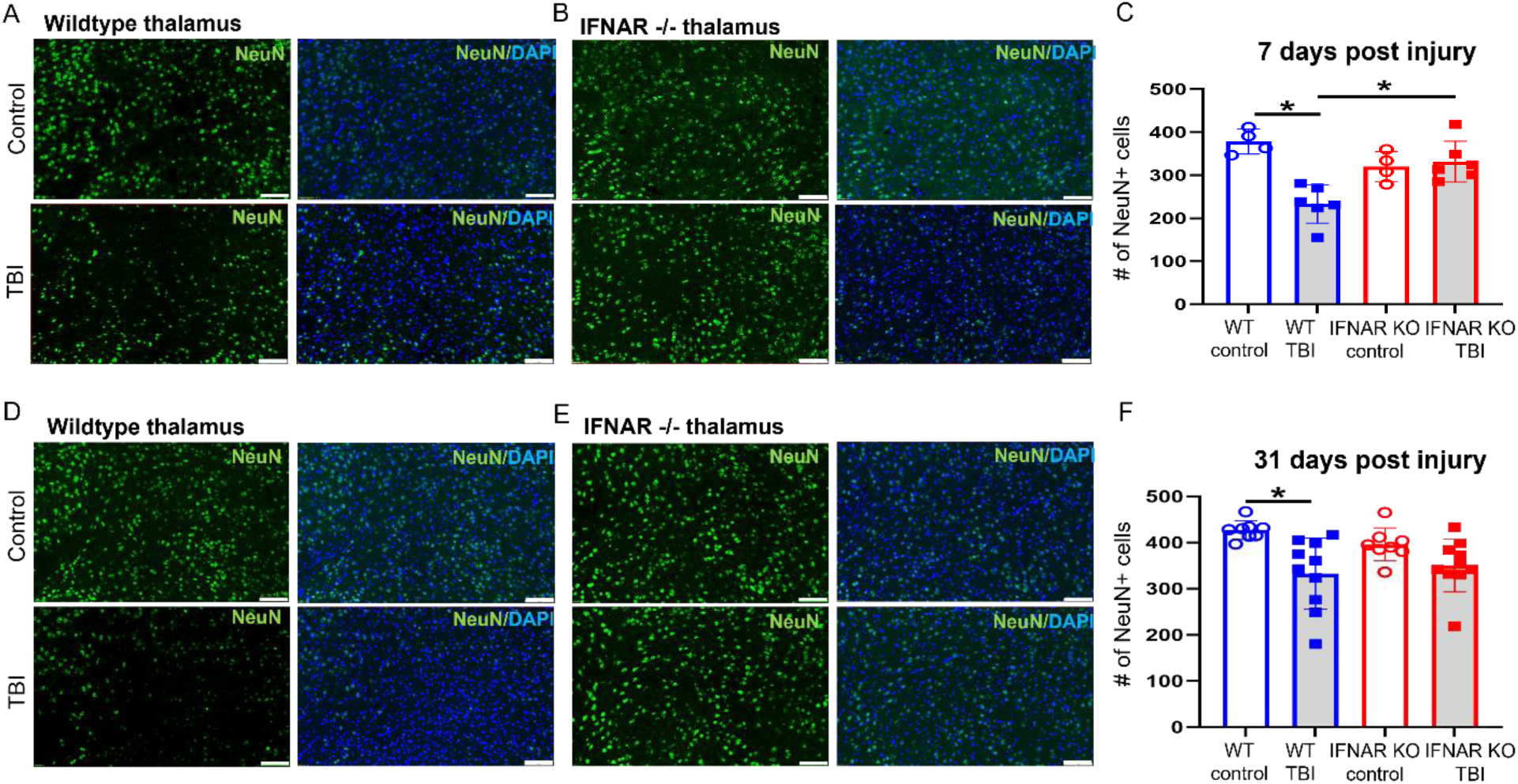
IFNAR deficiency reduces thalamic neuronal loss following TBI. Neuronal loss was quantified by IHC using NeuN (green) staining in WT and IFNAR KO subjects. Representative NeuN staining in the thalamus at 7 (**A** and **B**) and 31 days (**D** and **E**) following TBI or control. Scale bar=100 um. Quantification of NeuN+ cells in the thalamus (**C** and **F**). Error bars depict mean ± SEM. Statistical analysis with 2-way ANOVA with Fisher’s LSD for multiple comparisons. *p<0.05, n=4-8 control mice/group, n=6-10 TBI mice/group.

Clinically, traumatic brain injury often involves both focal and diffuse injuries to the brain (25, 26). Lateral FPI has the advantage of mirroring the mixed modality of brain injury that occurs in humans, making it possible to also evaluate how IFNAR deficiency affects white matter damage after TBI. Diffusion tensor imaging followed by tract-based spatial statistics was used to assess white matter disruption seven days following TBI. Fractional anisotropy, an indicator of axonal integrity, was significantly decreased in WT TBI mice compared to IFNAR KO TBI mice in perilesional white matter tracts (**Figure 9C**). Based on this evidence, we conclude that IFNAR deficiency reduced injury-induced disruption of the white matter at seven days post-TBI.

**Figure 9:**
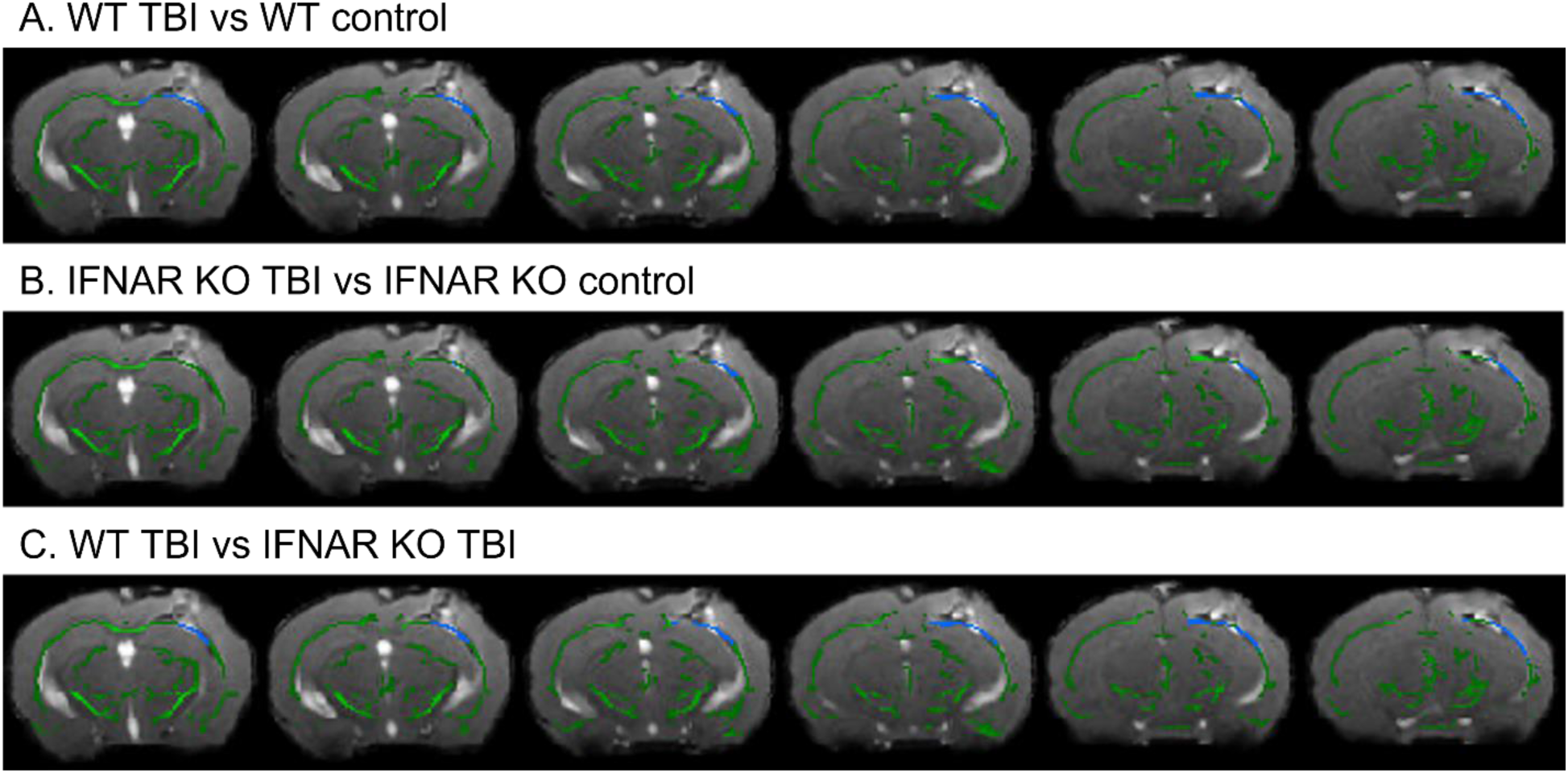
IFNAR deficiency reduces white matter disruption 7 days following TBI. Difference in white matter fiber tracts between WT and IFNAR KO TBI mice using diffusion tensor imaging. T-statistical maps of the lesion epicenter showing the decrease in fractional anisotropy (FA) in WT TBI vs WT control (**A**) and IFNAR KO TBI vs IFNAR KO control (**B**) at 7 DPI. The f-statistic of WT TBI vs IFNAR KO TBI (**C**) showing significantly larger decrease in FA in WT TBI compared to IFNAR KO TBI. Images are overlaid onto a representative TBI image with the mean FA skeleton shown in green. Areas of decreased FA are shown in blue. Distance from bregma in mm is shown along the bottom and distance from bregma shown below. n =15-17 mice/group.

### IFNAR deficiency ameliorates TBI-induced neurobehavioral dysfunction

To determine the role of interferon signaling on injury-induced neurologic dysfunction, we performed behavior testing in WT and IFNAR KO mice 2-3 weeks following TBI or in uninjured controls (**Figure 10A**). We first used the open field test to assess hyperactivity and anxiety phenotypes 14 days following TBI. As shown in **Figure 10B**, WT TBI mice displayed injury-induced hyperactivity with significantly increased total ambulation compared to WT control mice. In contrast, IFNAR KO mice did not develop injury-induced hyperactivity. To assess anxiety-like behavior, the total time in the center of the open field chamber was recorded for the duration of the trial. TBI did not increase anxiety-like behavior in either genotype with no difference from controls in time spent in the center of the open field (**Figure 10C**).

**Figure 10:**
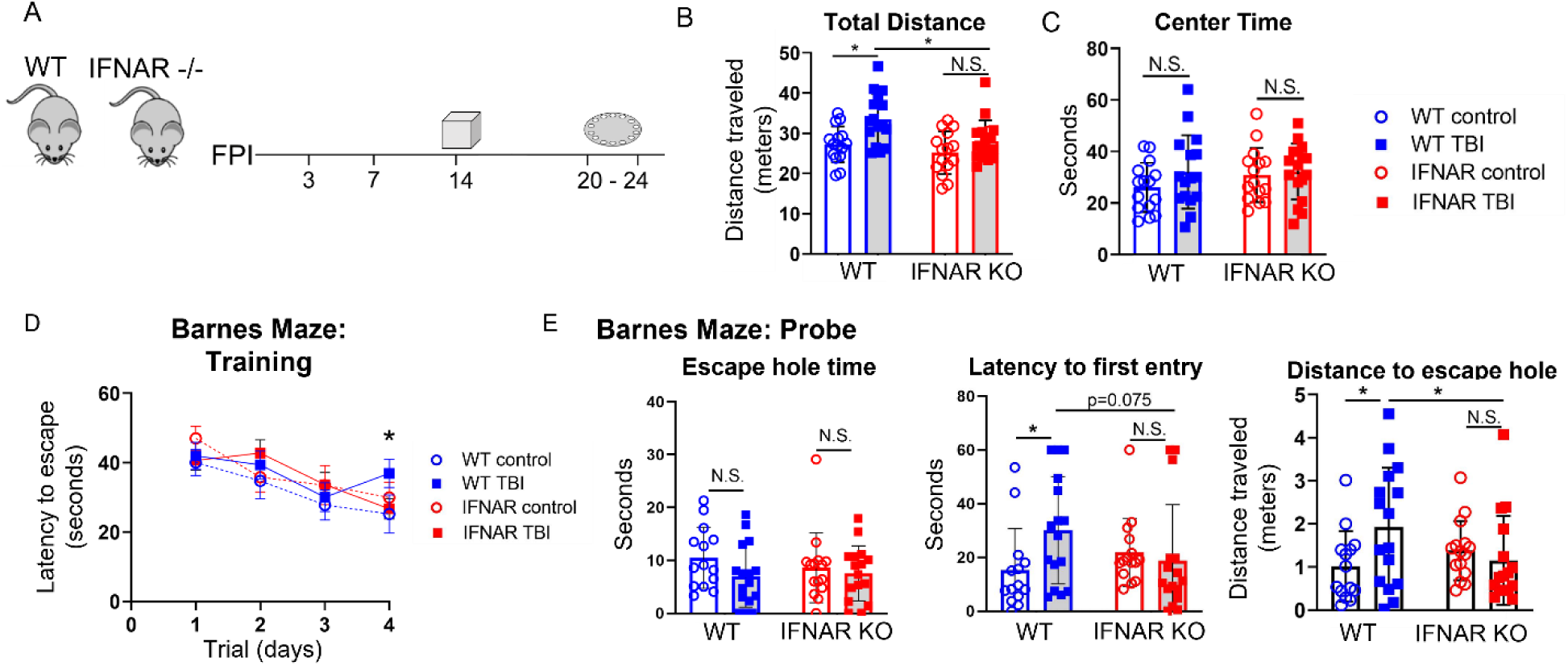
IFNAR deficiency rescues hyperactivity and improves spatial learning following TBI. (**A**) Graphic representation of the behavior paradigm. (**B** and **C**) 2 weeks following TBI, mice underwent open field testing. TBI induced hyperactivity in WT mice but not in IFNAR KO mice as indicated by increased total distance traveled (**B**) by WT TBI mice. There was no difference in the time spent in the center of the open field in TBI compared to uninjured control mice of either genotype (**C**). n=15-17 mice/group. 2-way ANOVA with Fisher’s LSD for multiple comparisons. *p<0.05. Barnes maze testing was performed 3 weeks post-TBI. (**D**) During the training phase, IFNAR KO TBI mice showed improved learning compared to WT mice exposed to TBI. (**E**) IFNAR KO mice exposed to TBI also showed improved memory during the probe trial. While there was no difference in time spent in the escape zone, IFNAR KO TBI mice had shorter latency and shorter distance traveled to first entry into the escape zone compared to WT TBI mice. n=15-17 mice/group. Error bars depict mean ± SD. Statistical analysis performed by 2-way RM ANOVA with Fisher’s LSD for multiple comparisons (**D**) or 2-way ANOVA with Fisher’s LSD for multiple comparisons (**E**). *p≤0.05

Next, we assessed spatial-based learning and memory using the Barnes maze. Testing was initiated on day 20 post-injury and extended through day 24. Spatial learning was evaluated during four days of Barnes maze training (**Figure 10D**). The main effect of time was significant. Post-hoc comparisons revealed IFNAR KO mice were protected from TBI-induced cognitive dysfunction. Only WT TBI mice performed worse than WT controls on day four of training. There was no significant difference between IFNAR KO TBI and control mice and a trending decrease in the latency to escape hole between IFNAR KO and WT injured mice. Overall this indicates that IFNAR KO mice had improved spatial learning compared to WT mice following TBI.

On the fifth day of Barnes maze testing, spatial memory was assessed during the probe trial (**Figure 10E**). There were no significant differences in total time spent around the escape hole. However, WT TBI mice took significantly longer than WT controls to enter the escape zone, consistent with spatial memory impairment. In contrast, IFNAR KO mice exhibited a trending decrease in time to first escape-zone entry compared to WT mice following TBI indicating improved spatial memory in the probe trial. The distance traveled en route to the first escape-zone entry was analyzed as a measure of path efficiency. WT TBI mice took a significantly less efficient path when compared to WT control mice. In contrast, IFNAR KO TBI mice showed no difference compared to uninjured controls. IFNAR KO TBI mice took a more efficient path compared to WT TBI mice. Collectively, mice with IFNAR deficiency were protected from injury-induced cognitive dysfunction including hyperactivity and spatial learning and memory deficits.

## Discussion

The IFN-I pathway is strongly upregulated across a wide spectrum of neurodegenerative diseases (12, 27–29). This robust response is the subject of increased study focused on the neurologic impact of IFN-I activation (30, 31). In TBI, we and others have demonstrated significant IFN-I activation both at the tissue level and in specific immune cell populations, including microglia and astrocytes (5, 9–11, 32–34). However, the effects of IFN-I signaling on the neuroimmune response to TBI and the mechanisms by which this response contributes to TBI-induced neuropathology are poorly understood. In this study, we demonstrate that IFN-I signaling drives expression of a specific subset of neuroimmune genes, alters microglial phenotype, and results in accumulation of microglia and peripheral leukocytes following TBI. Suppression of these neuroimmune responses by inactivation of the IFN-I receptor (IFNAR), was accompanied by reduced neuronal loss and white matter disruption and improved neurologic function following TBI. Our study highlights the therapeutic potential of IFNAR blockade and targeted immunomodulation following TBI.

The neuroimmune response to TBI is temporally dynamic and context-specific, with roles in both injury recovery and secondary neurotoxicity. Unravelling this complexity requires ongoing study to better understand the molecular signals that drive these diverse effects. Past work evaluating the impact of IFN-I signaling on the neuroimmune response to TBI has focused on small panels of genes, looking at their expression predominantly in the acute phase following TBI (10, 11, 32). To overcome this limitation, we performed multiplex gene analysis using the NanoString Neuroinflammatory panel, allowing broad unbiased screening of over 700 neuroinflammatory genes in WT and IFNAR KO mice following TBI. We also evaluated expression changes at both subacute and chronic timepoints. Somewhat unexpectedly, we found that both IFNAR-deficient TBI and WT TBI mice—relative to their uninjured control counterparts—had over 300 injury-induced DEGs at 7 DPI, with many DEGs overlapping between both genotypes. However, when comparing WT TBI and IFNAR-deficient TBI mice, we found that IFNAR deficiency potently suppresses a subset of 10 TBI-induced genes, previously identified as type I interferon-stimulated genes. This selective modulation differs from the findings of a prior study on Alzheimer’s disease whereby IFNAR deficiency resulted in suppression of both ISGs and complement pathway genes (12). Our findings are consistent with work in a mouse model of Parkinson’s disease; in this model, deficiency in STING, the upstream regulator of IFN-I, selectively modulated ISGs but did not prevent upregulated expression of complement and other inflammatory pathway genes (35). These disease-specific responses highlight the importance of context in driving neuroimmune regulation. In the case of TBI, the acute neurologic insult is more severe than in neurodegenerative diseases, and consequently, may trigger inflammation through activation of multiple, redundant mechanisms. It is important to note, however, that in both our study and the study of a Parkinson’s disease model, the selective prevention of IFN-I pathway activation was sufficient for improved neuropathology and neurobehavioral outcomes (35). In fact, the selective transcriptional alteration mediated by IFNAR deficiency may be a vital asset for development of highly specific IFN-I targeted therapies for TBI. As knowledge of the diverse states and functions of CNS immune cells has increased, there has been growing recognition of the importance of targeted vs broad manipulation of the immune response in the setting of neurologic disease. One cause of past failures of anti-inflammatory therapies following TBI may have been the broad immune cell suppression arising from these treatments, which would eliminate not only toxic but also reparative functions (36). Further study is needed to identify whether selective targeting of type I interferon signaling is sufficient and optimal for ameliorating harmful, dysregulated immune responses following TBI.

As the primary prenatally seeded immune cell in the CNS, microglia are important effector cells of the neuroimmune response following brain injury. While there is growing evidence for a pathologic role of IFN-I activated microglia in chronic neurodegenerative diseases, in TBI, the effects of IFN-I signaling on microglial accumulation and diverse microglial phenotypes are less understood (13, 27, 37, 38). One TBI study reported increased microglia accumulation in IFNAR-deficient mice 24 hours post-injury (11), while another demonstrated decreased acute microgliosis in mice deficient in STING, an upstream stimulator of IFN-I (32). As the acute microglial response may likely be necessary for debris clearance and other repair mechanisms, we aimed to study the effect of IFNAR signaling on the chronic microglial response. Additionally, we sought to address the impact of IFN-I on specific microglial functions. In our study, we found that IFNAR-deficient mice had significantly decreased regional microglial accumulation at 31 DPI, with the greatest effect found in the thalamus. We also demonstrated that type I IFN signaling drives microglial expression of MHC class I antigen processing and presentation factors following TBI. This altered microglial phenotype may be an important mechanism contributing to secondary injury following TBI. The requirement of microglial MHC class I antigen presentation for cytotoxic CD8+ T cell infiltration has been shown in CNS viral infection models (16). Furthermore, a prior study in TBI demonstrated that depletion of CD8+ T cells resulted in improved neurologic function following TBI (39). In our study, we showed that the decreased expression of microglial MHC class I molecules in IFNAR KO mice was associated with decreased brain accumulation of CD8+ T cells and was also associated with decreased neuronal loss, white matter injury, and neurobehavior dysfunction following TBI. Our ongoing studies will more directly evaluate the impact of IFN-I stimulated microglia on CD8+ T cell recruitment and subsequent neurotoxicity following TBI.

To modulate the IFN-I signaling pathway therapeutically, it is essential to know whether and how IFN-I impacts the multiple mechanisms of neuropathology involved in brain injury and disease. For example, in models of Alzheimer’s disease and frontotemporal dementia, IFN-I stimulated microglia cause pathologic synaptic loss (12, 13, 40). Additionally, in a Parkinson’s disease model, IFN-I pathway activation resulted in dopaminergic neuron loss (35). Following TBI, neuronal death, axonal injury, synaptic dysfunction, and white matter injury all can contribute to neurologic dysfunction. In this study, we demonstrated that thalamic neuron loss and white matter pathology were suppressed when IFN-I signaling was disrupted. This builds upon prior work that demonstrated a protective effect of type I IFN deficiency on hippocampal neurodegeneration, and cortical volume loss following TBI (10, 11). In our present study, loss of IFN-I signaling led to altered microglial phenotype and accumulation, coupled with a decrease in brain neutrophils, monocytes, and T cells following TBI. Previous studies using cell depletion strategies, have implicated activated monocytes, microglia, and T cells as contributors to chronic neuropathology following TBI (7, 41, 42). As such, it is likely that the neuroprotective effects of IFNAR deficiency are due to impacts on both centrally-and peripherally-derived immune cells—the result of altered immune cell crosstalk that influences neuroprotective and neurotoxic states. Our future work will seek to identify the specific ISGs that mediate TBI neuropathology. One molecule of particular promise is CXCL10. CXCL10 is known to be upregulated after TBI, and in our study, its expression was potently inhibited by IFNAR deficiency (33, 34). The exact role of CXCL10 in TBI neuropathology is unknown; however one study found that a single dose of intranasal CXCL10-siRNA prior to TBI resulted in decreased infiltration of Th1 cells (34). Another study in an experimental model of demyelinating disease found that CXCL10 deficiency resulted in reduced microglial activation and amelioration of chronic neurotoxicity (43). CXCL10 blockade following TBI warrants further investigation to dissect its role in TBI pathology. ZBP1 is another candidate molecule that is upregulated by TBI in WT mice, but whose transcription was prevented in our IFNAR KO TBI mice. While ZBP1 was initially identified as an innate immune sensor resulting in IFN-I upregulation, it is now also recognized to induce inflammatory cell death and thus warrants investigation as a potential effector molecule of neuropathology following TBI (44).

Our study has several limitations. To study the effect of IFNAR deficiency, we used a global IFNAR knockout mouse line. As with any global knockout mouse model, this carries the risk of altered neural and immunological development. Previous studies have shown decreased hippocampal synapse number and synaptic plasticity in healthy, IFNAR-deficient mice (45). Others have observed elevated levels of peripheral myeloid lineage cells in IFNAR-deficient mice (46). Additionally, while our studies did not find any significant behavioral differences when comparing IFNAR KO and wildtype controls, others have described spatial learning impairment in IFNAR-deficient mice (45). Future studies will utilize inducible IFNAR knockout and pharmacologic blockade to avoid the effects on development seen in the global knockout and to determine if IFN-I mediates differential effects on TBI pathogenesis at acute, subacute, and chronic timepoints. We will also use cell-type-specific IFNAR-deficient KO models to dissect the cell-specific contributions to type I interferon-driven neuropathology following TBI. Finally, a limitation in this study is the exclusive use of male mice. It is well established that sex is a relevant biologic variable after TBI (47, 48). Others have demonstrated sex differences in the neuroimmune response to TBI, including decreased inflammatory activation and reduced infiltration of myeloid cells in female mice (49). Further study is needed evaluate IFN-I signaling in both sexes.

In summary, this study shows that IFN-I signaling is widespread and persistently upregulated following TBI. The rapid and persistent upregulation of the IFN-I pathway reveals a broad therapeutic window for intervention. IFNAR deficiency results in selective modulation of both the central and peripheral neuroimmune response. This includes reduced neuroinflammatory gene expression, decreased regional microglial accumulation and transcription of antigen presentation genes, and decreased monocyte and T cell accumulation. The neuroimmune modulation of IFNAR-deficient mice is associated with reduced neuronal loss, reduced white matter disruption, and amelioration of TBI-induced neurobehavioral dysfunction. Overall, blockade of type I IFN signaling after a traumatic brain injury is a promising approach for developing immune-modulating therapies that may confer neuroprotection following TBI.

## Methods

### Animals

All studies were conducted on adult, 2-6 month old, male C57BL/6J (#000664) or global IFNAR1-deficient mice (#028288; B6(Cg)-*Ifnar1^tm1.2Ees^*/J, *Ifnar1* null allele) purchased from Jackson Laboratory. Average weight on day of craniectomy was 26.0 ± 3.6 g. Mice were housed in the Animal Care Facility at the University of Iowa (Iowa City, IA) under a 12-h light-dark cycle with *ad libitum* access to food and water. After craniectomy and fluid percussion injury (FPI) or non-surgical control, all mice remained singly caged. All procedures performed in this study were in accord with protocols approved by the Institutional Animal Care and Use Committee at the University of Iowa.

### Fluid percussion injury

Lateral FPI was performed as previously described (50, 51). On the day preceding injury, mice underwent craniectomy. Animals were anesthetized with ketamine/xylazine (87 mg/kg ketamine and 12 mg/kg xylazine) via intraperitoneal injection. The head was then mounted in a stereotaxic frame, and a midline incision of the scalp was made for reflection of the skin and exposure of underlying skull. A 3-mm OD handheld trephine (University of Pennsylvania Machine Shop) was used for craniectomy on the left parietal skull bone centered between lambda and bregma sutures and between lateral skull edge and sagittal suture. A modified Luer-Lock hub was placed surrounding the craniectomy site and secured with cyanoacrylate glue (Loctite 760355). The hub was further secured with methyl-methacrylate dental cement (Jet Acrylic Liquid mixed with Perm Reline/Repair Resin) surrounding the bottom portion of the hub. The hub was filled with sterile saline and closed with a sterile intravenous cap to prevent dural exposure to the environment.

The following day, mice underwent FPI. The pendulum angle of the FPI device was adjusted before use on each experimental group to achieve a peak pressure between 1.3 and 1.5 atmospheres (atm) when triggered against capped intravenous tubing. For experiments in this study, the pendulum angle varied between 10.8 and 11.8 degrees. Mice received 3% inhaled isoflurane in an induction chamber before being transferred to a nose cone, where the intravenous cap was removed and any air bubbles in the hub were eliminated.

Once deeply anesthetized, mice were connected to the FPI device via 20-inch IV tubing and placed on their right side. The pendulum was released, generating a brief fluid pulse against the exposed dura. A Tektronix digital oscilloscope (TDS460A) was used to measure the duration and peak pressure of the fluid pulse. After injury, mice were placed on their backs, and their righting time was measured as an indicator of injury severity. After righting, mice were re-anesthetized with isoflurane, the Luer-Lock hub was removed, and the skin incision was sutured closed. After skin closure, anesthesia was discontinued, and animals were placed in a heated cage until recovered and ambulatory. Given we were interested in studying moderate to severe traumatic brain injury, mice were included only if the duration of the righting reflex was >4 min (52–54). Across all studies, the average righting time ± SD was 380 ± 79 sec, which corresponded to an average peak pressure delivered of 1.4 ± 0.05 ATM. For Figure 1, mice received sham injury where they underwent identical treatment through connection to the FPI device but were disconnected without triggering of the FPI device. For the data presented in all the other figures, non-surgical controls were used; these mice received anesthesia and analgesia but did not undergo craniectomy.

### RNA-scope and Immunohistochemistry

Mice were anesthetized with ketamine/xylazine and perfused with ice-cold saline followed by 4% paraformaldehyde (PFA) 7 or 31 days after TBI or control (no TBI).

Dissected brains were post-fixed in 4% PFA at 4 °C overnight, then cryoprotected in a 30% sucrose solution until sinking. Brains were embedded in optimal cutting temperature (OCT) compound by the University of Iowa Central Microscopy Research Facility, and 10-μm coronal sections were prepared. Frozen sections from the injury epicenter were dried at 40 °C for 30 min prior to staining.

The RNAscope Multiplex Fluorescent detection Kit v2 was used per manufacturer’s instructions to stain for mRNA expression of *Cxcl10* (ACD, 408921) or *H2-K1* (ACD, 1049831-C1). For all immunohistochemistry, slides were placed in a blocking/extraction solution (0.5% Triton X-100 and 10% goat serum in 1× PBS) for 1 h at room temperature. After blocking, tissues were incubated overnight at 4 °C in rabbit anti-IBA1 (1:200; Wako Chemicals 019-17941), rabbit anti-CD8α (1:500; Abcam 217344), or rabbit anti-NeuN (1:200; Abcam 177487) primary antibody diluted in blocking/extraction solution. Alexa Fluor-488 or -568 conjugated goat anti-rabbit secondary antibody (Life Technologies) was applied at a 1:500 dilution for 1 h at room temperature. Fluorescently stained tissue slices were imaged using a slide-scanning microscope (Olympus VS120). Regions of interest were demarcated using OlyVIA software (Olympus). ImageJ was used to perform proportional area analysis (IBA1+) and quantification of NeuN+ cells.

### RNA Isolation and NanoString Gene Expression Analysis

7 or 31 days after control or TBI, mice were euthanized with isoflurane prior to decapitation and removal of the brains. Brain tissue was dissected, collected by region, and snap-frozen in liquid nitrogen. Total RNA was extracted from control or TBI brain regions using TRIzol (Invitrogen, Carlsbad, CA) as per the manufacturer’s instructions. RNA quantity and quality were evaluated with the Agilent 2100 Bioanalyzer. Gene expression was quantified using the NanoString nCounter Neuroinflammation panel. Data was normalized using nSolver software with a background threshold count value of 20. Housekeeping genes *Gusb* and *Asb10* were flagged due to differential expression across treatment groups and were excluded. The geometric mean was used to compute the normalization factor. Differential gene expression analysis was carried out with DEseq2. Genes that featured a log2fold change >0.5 and an adjusted p value < 0.05 were considered to be differentially expressed genes (DEGs).

### Quantitative real-time PCR

First-strand complementary DNA (cDNA) was synthesized with SuperScript III reverse transcriptase (Invitrogen). Amplified cDNAs were diluted 1:15 in ultra-pure water and subjected to real-time polymerase chain reaction (PCR) on an Applied Biosystems Model 7900HT with TaqMan Universal PCR Mastermix (Applied Biosystems, Foster City, CA) and the following Taqman probes: *Irf7* (Mm00516793), *Rsad2* (Mm00491265), *Slfn8* (Mm00824405), *Ddx58* (Mm01216853), *Ly6a* (Mm00726565), *Ifih1* (Mm00459183), *Ifitm3* (Mm00847057), *Zbp1* (Mm01247052), *Cxcl10* (Mm00445235) *H2-K1* (Mm01612247), *β2m* (Mm00437762), *Tap1* (Mm00443188), and *Gapdh* (Mm99999915). PCR reactions were conducted as follows: 2 min at 50°C, 10 min at 95°C, followed by 40 cycles for amplification at 95°C for 15 sec and 60°C for 60 sec. Biologic samples were run in duplicate or triplicate. Genes of interest were normalized to endogenous control *Gapdh*. Data were analyzed using the comparative cycle threshold method, and results are expressed as fold difference from WT controls.

### Open Field

Hyperactivity and anxiety-like behavior were assessed using the open field test. The open field apparatus (SD Instruments) consisted of a single enclosure 20 x 20 x 15 inches (0.51 x 0.51 x 0.38 meters) with a center area of 10 x 10 inches (0.25 x 0.25 meters). 14 days after control treatment or TBI, open field testing was performed. During testing, mice were placed individually in the center of the open field enclosure, and they moved freely for 5 minutes while their locomotor activity was recorded by a video camera mounted above the apparatus. Overall ambulatory movement was quantified by the total body distance traveled during the trial. Total distance traveled and time spent in the center were reported automatically by the ANY-Maze Video Tracking System (Stoelting, IL, USA). After each test, the apparatus was thoroughly cleaned with 75% ethanol.

### Barnes Maze

Cognitive function was assessed using the Barnes maze (SD Instruments). The Barnes maze was a white circular table 36 inches in diameter with 20 holes, 2 inches in diameter, evenly spaced around the perimeter. The table was brightly lit and open, motivating the test subjects to learn the location of the dark escape box located under one of the 20 holes. The maze was enclosed with four different visual cues hung on each wall surrounding the table. ANY-maze video tracking was used for data collection.

Three weeks after control or TBI, acquisition trials were conducted (four trials per day) for 4 days, during which an escape box was placed under the target hole. Each trial ended when the mouse entered the target hole or after 80 seconds had elapsed. Mice that did not locate the escape box were guided to the target hole where they entered the escape box. All mice were allowed to remain in the escape box for 15 sec before removal from the apparatus. Average latency to the escape hole was recorded for each acquisition day. On day 5 of Barnes maze testing, a probe trial was conducted to assess memory. The escape box was removed from under the target hole and mice were placed in the maze for 60 sec. Each mouse underwent one probe trial, during which the time spent in a 2-cm-diameter zone around the target hole, the latency to first escape-zone entrance, and the distance to first escape-zone entrance was recorded. Mice that displayed > 20 seconds without video tracking (2 subjects) on the probe trial were excluded from probe trial data analysis.

### Flow Cytometry

Three or ten days after TBI, mice were euthanized and their brains were removed. Mononuclear cells were isolated by digesting brain tissue in CollagenaseD/DNase (Sigma, 1108886601, D4513) for 45 min at 37 °C, dissociating through a 70-μm filter and isolating the cells at the interface between a 70% and 37% Percoll gradient (GE, 17-0891-01) after a 20 min, 25 °C spin at 2,000 r.p.m. Single-cell suspensions were plated and stained for 20 min at 4 °C with a combination of fluorescently labeled antibodies that are specific for surface markers. Cells were enumerated with counting beads (ThermoFisher, C36950) and then analyzed by flow cytometry. All data for samples were acquired using an LSRFortessa^TM^ Cell Analyzer (BD Bioscience) and analyzed using FlowJo software, v.10.8.1 (FlowJo LLC).

### Flow Antibodies

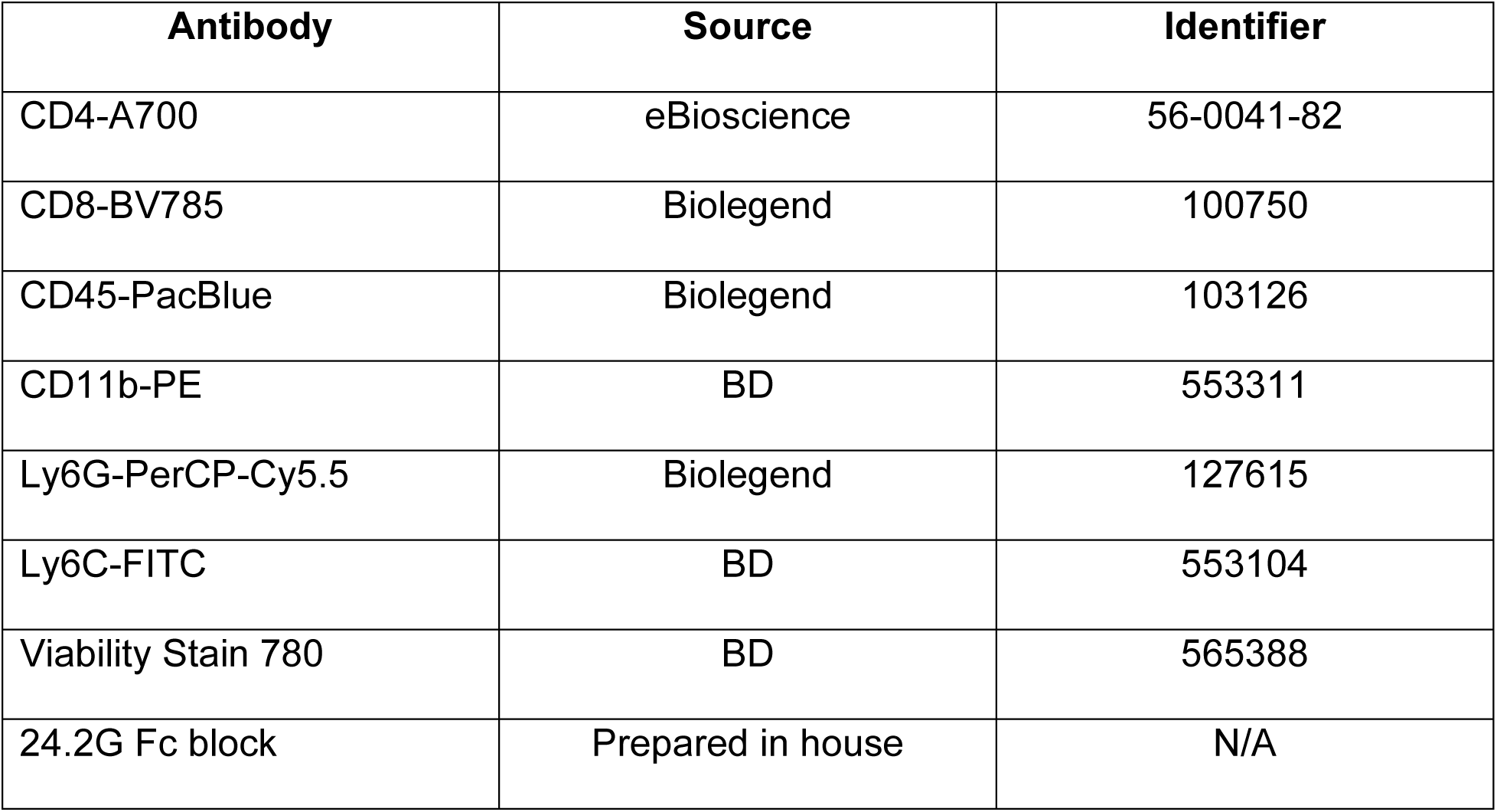

**Flow Antibodies:**
| **Antibody** | **Source** | **Identifier** |
| --- | --- | --- |
| CD4-A700 | eBioscience | 56-0041-82 |
| CD8-BV785 | Biolegend | 100750 |
| CD45-PacBlue | Biolegend | 103126 |
| CD11b-PE | BD | 553311 |
| Ly6G-PerCP-Cy5.5 | Biolegend | 127615 |
| Ly6C-FITC | BD | 553104 |
| Viability Stain 780 | BD | 565388 |
| 24.2G Fc block | Prepared in house | N/A |

### MRI

MRI images were obtained at baseline and 7 days post injury using the 7.0T GE 901 Discovery MRI Small Animal Scanner housed at the University of Iowa MR Research Facility. Mice were placed in an induction chamber with isoflurane (3%) until anesthetized, then each mouse was transferred to the scanner and placed in a specially designed imaging tray with a nose cone for continued anesthesia (1-1.5%). Diffusion tensor images (DTI) were obtained, and each scan lasted approximately 30 minutes. DTI was used to evaluate the microstructural integrity of the white matter including evaluation of the fractional anisotropy (FA) using DITFIT followed by tract-based spatial statistics (TBSS) (55). T-statistics were performed on fiber tract skeletons from WT and IFNAR KO animal to look for changes following TBI compared to baseline scans. The results of those t-statistics were the input for the calculation of the f-statistic to determine if any group differed from the mean.

## Acknowledgements

EAN received support from the Iowa Neuroscience Institute and the Carver Trust. T.N.-J. was supported by the Andrew H. Woods Professorship INI Carver

