## Supplementary figures and images for "Selective neuroimmune modulation by type I interferon drives neuropathology and neurologic dysfunction following traumatic brain injury"

### Supplemental Figure 1

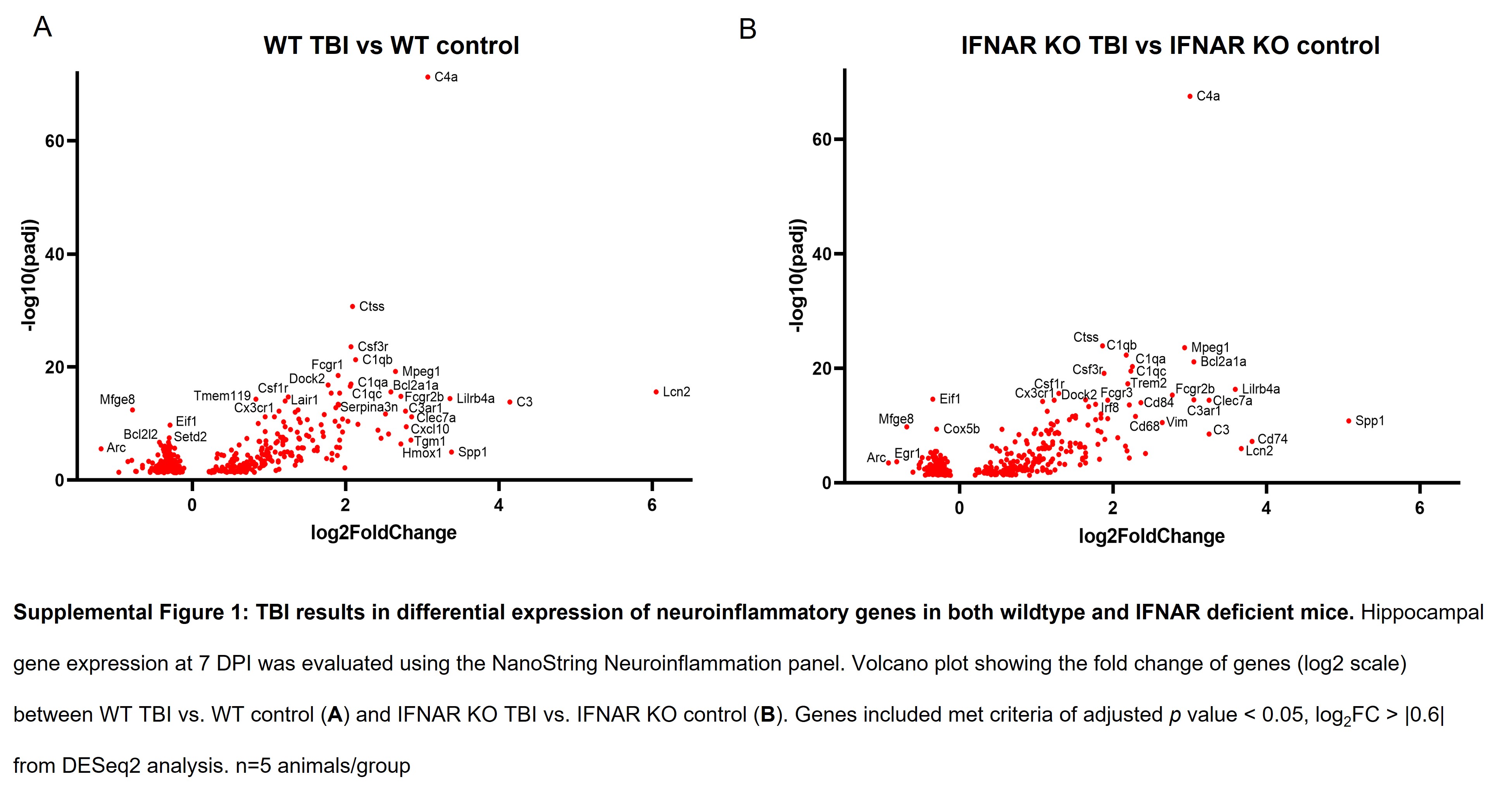
